# A demographic framework for assessing population vulnerability to contrasting perturbation regimes

**DOI:** 10.64898/2026.05.22.727074

**Authors:** Àlex Giménez-Romero, Daniel Oro, Daniel F. Doak, Mª Begoña García, Meritxell Genovart

## Abstract

Environmental change affects demographic rates through perturbations that differ in magnitude, duration, and frequency, yet their consequences for population vulnerability, i.e., potential population reduction, remain only partly understood. Here, we develop a general demographic framework that unifies pulse and press perturbations to better understand how life-history strategy shapes population declines across the fast–slow continuum. Using matrix population models for 12 plant and animal species with diverse generation time and life history strategies, we simulated perturbations acting independently on adult survival, juvenile survival, and fecundity, and measured their demographic consequences over comparable life-history timescales. We then integrated impacts across perturbation regimes to derive a novel comparative vulnerability metric and related this metric to species’ life-history descriptors. Across taxa, perturbations to adult survival consistently produced the strongest demographic impacts, with vulnerability increasing markedly towards slower life histories. Juvenile survival emerged as the main axis of demographic differentiation among species, whereas the effects of perturbations on fecundity were weaker and comparatively homogeneous across the continuum. Generation time strongly predicted vulnerability to survival perturbations, but not to reproductive output. Consistent with previous theoretical and empirical work, our results show that vulnerability is not a fixed species property, but an emergent outcome of the interaction between the perturbed vital rate, the temporal structure of environmental forcing, and the underlying life-history strategy. Importantly, as the vulnerability metric can be compared both across populations under a given perturbation regime and within populations across perturbation types and demographic targets, the framework also provides a basis for stage-specific and regime-specific management.

## 1 Introduction

Environmental variability is an inherent feature of natural systems that exposes populations to fluctuations in the conditions governing survival, development, and reproduction [1, 2]. These fluctuations can generate perturbations in demographic rates that vary in magnitude, duration, and temporal structure [3–6]. Understanding how populations are affected by such perturbations is central to ecology, as these impacts largely determine whether populations persist, decline, or recover following environmental change [7–10]. Species exhibit substantial diversity in life-history strategies, differing in how they allocate resources to survival, development, and reproduction, in the timing of reproductive events, and in overall life-cycle length[11]. These life-history differences also shapes the sensitivity of demographic rates to perturbations, such that species differ in which vital rates are most affected when the environment changes [12, 13]. While substantial progress has been made in understanding how populations are affected by demographic perturbations across life-history strategies, further work is needed to refine how vulnerability, defined here as the proportional reduction in population size caused by perturbations, is quantified and assessed, particularly through complementary demographic perspectives. Such advances will improve understanding of population dynamics under variable environments and strengthen our ability to manage populations more effectively [14, 15].

Environmental perturbations affecting demographic rates can differ not only in their intensity, but also in how they unfold over time. They may be punctual in time or sustained pressures continuously perturbing the system over multiple time steps [16, 17]. These contrasting temporal regimes are often conceptualized as pulse or press [16], and are known to shape population responses in distinct ways [18, 19]. For example, brief perturbations may be partially buffered by compensatory dynamics or appear less evident due to demographic inertia, whereas prolonged or gradually intensifying perturbations can lead to cumulative effects that overwhelm recovery mechanisms [20, 21]. Despite this, the temporal structure of perturbations is often simplified or ignored, limiting our ability to compare impacts on populations or species, across different forms of environmental variability [22, 23].

Much of the recent work on demographic responses to disturbances has focused on how populations respond to pulse perturbations on some of their vital rates [24–26]. By contrast, press perturbations and other temporally extended forms of demographic forcing remain comparatively underexplored within a unified comparative framework [27–29]. This is a major limitation, because many real environmental drivers are not well described as isolated shocks. Climate anomalies, habitat degradation, harvesting, disease outbreaks, or chronic resource limitation often act over multiple time steps, recur before populations fully recover, or combine moderate intensity with long duration [19, 30–32]. Under these conditions, demographic impact is expected to depend not only on which vital rate is perturbed, but also on how perturbation magnitude, duration, and recurrence interact with the species’ life-history pace. Approaches that treat these dimensions separately make it difficult to compare vulnerability across perturbation regimes and across taxa. A general framework that places pulse and press perturbations within the same demographic space is therefore needed to evaluate how life-history strategy modulates population vulnerability under environmental forcing.

From a methodological point of view, structured population models provide a natural framework for formalizing how perturbations affect demographic processes, linking changes in individual-level vital rates to population-level outcomes [4]. Matrix population models (MPMs), in particular, describe population dynamics through a projection matrix whose elements encode transitions among life stages and reproductive output [13, 33, 34]. Changes in specific matrix elements can therefore be interpreted directly as perturbations acting on well-defined demographic processes. Related approaches such as integral projection models (IPMs) are also widely used, particularly for size-structured populations, and are based on the same demographic principles and commonly analyzed using equivalent matrix-based tools [35, 36]. Because these models explicitly represent life-history structure, they have become standard tools for exploring how life-history strategies shape population dynamics across taxa [13, 37, 38].

Previous works have largely focused on modeling pulse perturbations acting on specific vital rates as discrete, single-step changes in population structure [24–26]. In this context, perturbations are typically represented as instantaneous changes in the population state vector, and population impacts are quantified through transient dynamics such as amplification, reactivity, or rates of return to equilibrium [25]. This line of research has yielded important insights into how life-history traits shape short-term population responses and recovery following sudden perturbations [39]. However, these approaches generally assume that the underlying demographic parameters remain constant through time. This limits the applicability of these frameworks to many ecological situations in which environmental change acts on different demographic rates, such as survival or reproduction, and does so over sustained periods or with non-instantaneous temporal structure [4, 6, 26].

Here, we develop a framework that systematically explores how perturbations differing in magnitude, duration, and frequency affect distinct vital rates across species spanning a broad range of life-history strategies. This enables a comparative assessment of how life-history variation interacts with disturbance characteristics to shape demographic vulnerability across taxa. The selected species primarily derive from long-term demographic studies with robust parameter estimates, complemented by additional species from COMADRE [15] to ensure broad coverage of generation times. Using matrix population models, we simulate population dynamics under perturbations independently affecting adult survival, juvenile survival, or fecundity, while treating pulse and press regimes as part of a single continuous perturbation space. By jointly varying perturbation intensity, duration, and recurrence, we quantify the resulting demographic impact and derive an integrated measure of vulnerability over ecologically relevant timescales. We then relate these responses to a comprehensive suite of life-history descriptors derived from each MPM, allowing us to identify the demographic traits and strategies that best predict vulnerability across perturbation types. Our approach provides a general comparative framework for assessing species vulnerability and for understanding how variation in life-history strategy shapes the demographic consequences of environmental change.

## 2 Methods

### 2.1 Population modeling framework

We evaluated the effect of different perturbation regimes to the demographic parameters of several plant and animal species with different life history strategies using matrix population models (MPMs). For each species, we considered a stage-structured model of the form

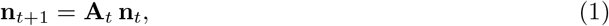

where **n**_*t*_ is the population vector at time *t* and **A**_*t*_ is the projection matrix governing transitions between life-history stages. The projection matrix includes stage-specific survival and transition probabilities as well as fecundity terms. All simulations were run for 3 generations (*t*_*max*_ = 3 *T*_*R*_0, see Methods below), which we consider an ecologically relevant timescale providing sufficient time for impacts to emerge.

### 2.2 Study species and demographic data

To enable a comparative analysis of perturbation effects across taxa, we selected 12 species with robust demographic data representing a broad spectrum of life-history strategies along the fast– slow continuum. The dataset encompasses both plant and animal species, including mammals (*Mus musculus, Elephas maximus*), birds (*Cyanistes caeruleus, Ichthyaetus audouinii*), amphibians (*Alytes muletensis*), reptiles (*Dermochelys coriacea*), fishes (*Poecilia reticulata*), herbaceous plants (*Mimulus cardinalis, Mimulus lewisii*), shrubs (*Hibiscus meyeri*), and trees (*Pinus nigra, Quercus rugosa*). Most species derive from our own long-term demographic monitoring programs [40–44], providing well-validated demographic parameter estimates, while additional species were obtained from the COMPADRE/COMADRE database [15, 38] to further expand coverage across generation times, life-history strategies, and taxonomic groups. Together, the selected species span a wide range of generation times, from fast-lived organisms with short generation times of a few years to long-lived species whose generation times extend over several decades. This broad comparative coverage allows us to examine whether vulnerability to perturbations varies systematically with life-history traits. Each species was represented by a stage-structured matrix population model (MPM) including survival, stage-transition, and fecundity processes. All employed MPMs with their corresponding specific references can be found in Supplementary Section 1.

### 2.3 Life-history descriptors

To characterize the variability of life-history strategies, we computed a set of standard demographic descriptors derived from the unperturbed projection matrix for each species. These descriptors summarize reproductive output, timing of reproduction, survival patterns, age structure, and the dispersion of demographic events across the life cycle. All quantities were computed using the MatPopMod Python library [45], which provides a unified framework for analyzing stage-structured population models and associated life tables. The complete list of species and their associated life-history descriptors are shown in Supplementary Section 2.

#### 2.3.1 Reproductive output

Reproductive output was quantified using the net reproductive rate, *R*_0_, defined as the expected number of offspring produced by an individual over its lifetime. In matrix population models, *R*_0_ can be expressed as

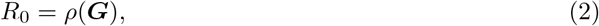

where *ρ* denotes the dominant eigenvalue and **G** = ***F*** (***I*** −***S***)^*−*1^ is the next generation matrix, derived from the fecundity and survival–transition sub-matrices, ***F*** and **S**, respectively [13].

In addition, we computed the cohort-based net reproductive rate as the average number of offspring that a newborn is expected to produce over its lifetime in the stable population, which is given by,

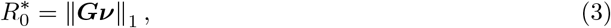

where ***v*** is a vector containing the fraction of newborns that are born in each class when the population is at its age stable distribution.

#### 2.3.2 Generation time

To characterize the temporal scale of reproduction, we computed several complementary measures of generation time. The generation time associated with *R*_0_, denoted 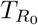, was defined as

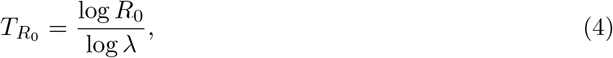

where *λ* is the population growth rate.

We also computed the mean age of mothers in the stable population, *T*_*a*_,

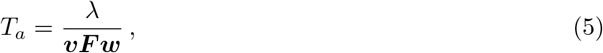

where ***v*** and ***w*** are the left and right eigenvectors of the projection matrix, respectively.

Furthermore, we computed the cohort generation time, *µ*_1_, which computes the mean age of the mothers when considering all offspring produced by a cohort,

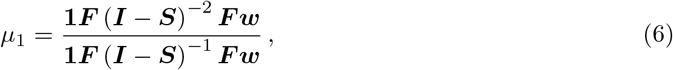

where **1** is a vector of ones, and a variant of this measure where births are weighted by the cohort reproductive values ***v***_*G*_ instead of equally,

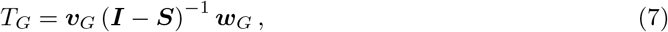

where ***v***_*G*_ and ***w***_*G*_ are the left and right eigenvalues of the next generation matrix, respectively.

Finally, we also estimated the mean age of reproduction numerically from the stable age distribution by sampling reproductive events until convergence.

#### 2.3.3 Life expectancy

Survival patterns were summarized using life expectancy, defined as the expected number of projection intervals that an individual born in the stable population lives, and was computed as,

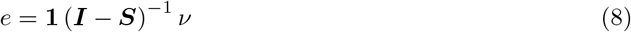

We also numerically computed the life expectancy conditioned on reproduction, *e*_*r*_, based on Markov chains (see [45]).

#### 2.3.4 Age structure

Population age structure was characterized by the mean age of individuals in the stable age distribution, computed as,

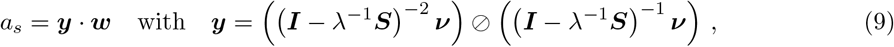

where ⊘ corresponds to the entry-wise division operation.

#### 2.3.5 Entropy

We quantified the dispersion of demographic events across the life cycle using entropy-based measures. Population entropy was computed from a genealogical Markov chain, ***P***, as

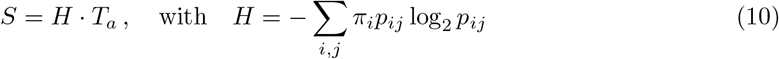

where *H* is the entropy rate of the model.

In addition, we also computed the life-table entropy, also known as Keyfitz’s entropy, which summarizes the distribution of mortality across ages based solely on the survival schedule,

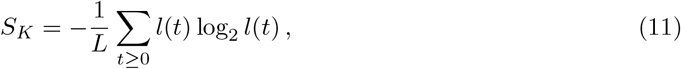

where *l*(*t*) is the probability of reaching age t.

### 2.4 Demographic perturbations

Perturbations were implemented as targeted reductions in selected elements of the projection matrix. We considered three classes of demographic parameters: (i) adult survival, defined as the survival probability of reproductive or sexually mature stages; (ii) juvenile survival, defined as the survival probability of pre-reproductive stages; and (iii) fecundity, defined as the number of offspring produced per reproductive individual. Perturbations were applied as multiplicative reductions of the corresponding matrix elements. A perturbation of magnitude *m* therefore replaced the unperturbed value *x* by (1 −*m*)*x*, where *m* represents a proportional reduction. Magnitude values ranged from 0 to 1, corresponding to proportional decreases in the targeted demographic parameter.

To capture contrasting perturbation regimes, we specified their temporal structure through duration *d* and frequency *v*. Duration corresponds to the number of consecutive time steps during which the demographic parameters are modified. A duration of one time step represents a pulse perturbation, whereas durations greater than one represent press perturbations. Durations ranged from 0 time steps to the number of steps corresponding to one generation. Frequency determines the interval between repeated perturbation events; a frequency of *v* implies that the perturbation is imposed every *v* time steps over the simulation window. For each combination of magnitude, duration, and frequency, the projection matrix **A**_*t*_ was updated accordingly at the specified time steps.

To quantify the demographic consequences of each perturbation regime, we evaluated the population size at the end of each 3-generation simulation. To isolate the persistent demographic consequences of each perturbation regime from short-term transient fluctuations, once the perturbation sequence had ended we projected the population for four additional generations using the unperturbed matrix. Final population sizes were then measured after this period. This procedure allowed us to quantify the net demographic impact of the perturbation regime while minimizing the influence of transient effects induced by the perturbations.

All perturbation durations, frequencies, and simulation horizons were defined relative to generation time rather than calendar time. This choice allows comparisons across species with markedly different life histories on a common demographic scale, so that a perturbation is interpreted relative to the pace of the life cycle rather than its absolute duration in years. The resulting framework is therefore most directly suited to comparative analyses of demographic vulnerability across species, rather than to management questions framed over fixed calendar horizons. However, one can use our framework without applying this rescaling to answer specific management-related questions.

### 2.5 Global impact and population vulnerability

To assess the global impact to perturbations, we used the estimated population size at the end of each 3-generation simulation and evaluated the percent population reduction (*ρ*). For a given perturbation regime (*m, d, v*), we defined vulnerability as the percent population reduction,

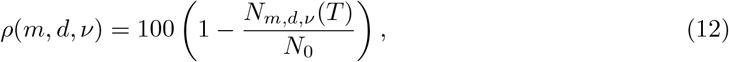

where *N*_*m,d,v*_(*T*) is the final population size after *T* time steps under the perturbed dynamics, and *N*_0_ is the initial population size, which would correspond to the final population size in the absence of perturbation. Thus, *ρ* = 0 indicates no demographic effect, whereas *ρ* = 100 corresponds to complete population extinction relative to the unperturbed trajectory. This definition allowed us to characterize then the effect of perturbations over the full perturbation space, Ω = { *m, d, v*}, spanned by all simulated values of perturbation magnitude, duration, and frequency. This metric is independent of absolute population size.

In contrast to classical sensitivity or elasticity analyses, which describe the effect of small changes in vital rates on asymptotic population growth, our metric explicitly incorporates the temporal dimension of perturbation. As a result, *ρ*(*m, d, v*) depends not only on which demographic parameter is affected, but also on the magnitude, duration, and frequency of the perturbation. This makes it possible to compare the demographic impact of different temporal regimes, from short pulse perturbations to sustained or recurrent press-like perturbations.

To synthesize the overall vulnerability of a population into a single quantity, we then averaged the percent reduction over the entire perturbation space,

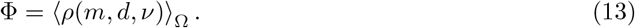

Equivalently, in discrete form,

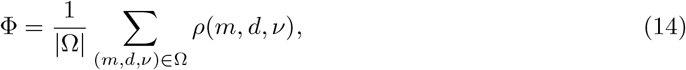

where |Ω| denotes the total number of simulated perturbation regimes. Thus, Φ measures the mean demographic impact of perturbations after jointly integrating over their magnitude, duration, and frequency. Higher values of Φ indicate populations that are, on average, more vulnerable across the perturbation regimes explored, whereas lower values correspond to populations that remain comparatively unaffected throughout the perturbation space.

### 2.6 Relating life-history descriptors to population vulnerability

To relate differences in species’ vulnerability to life-history variation, we examined how the vulnerability metric Φ co-varied with the descriptors defined in Section 2.3 across the 12 study species. These analyses were conducted separately for perturbations affecting adult survival, juvenile survival, and fecundity, and also for a fourth scenario in which all three demographic parameter classes were perturbed simultaneously.

For each perturbation affecting different demographic parameters, we fitted a separate univariate linear regression using Φ as the response variable and each life-history descriptor as the predictor. Because several descriptors span multiple orders of magnitude across species, we considered both raw and log-transformed predictor values when appropriate and retained the transformation that provided the strongest linear fit. Descriptor importance was quantified using the coefficient of determination (*R*^2^), and statistical significance was assessed from the slope term of each regression, using *p <* 0.05 as the significance threshold. Given the limited number of species and the strong collinearity expected among many life-history descriptors, these analyses were intended as a comparative screening of predictor strength rather than as a multivariate model-selection exercise.

### 2.7 Elasticity analysis

To assess the additional contribution of our integrated vulnerability metric compared to the classical asymptotic perturbation measures, we computed the elasticity of the asymptotic population growth rate, *λ*, to changes in the elements of each projection matrix. For an unperturbed matrix element, *a*_*ij*_, elasticity is defined as

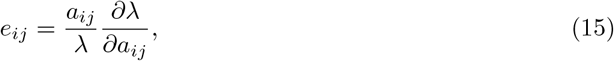

which quantifies the proportional change in *λ* resulting from a proportional change in *a*_*ij*_ [12, 13]. Elasticities were calculated from the dominant left and right eigenvectors of the projection matrix following standard matrix population model theory,

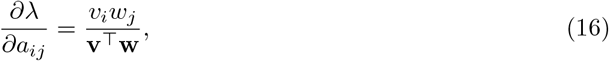

where **v** and **w** are the dominant left and right eigenvectors of the projection matrix, respectively.

To obtain quantities comparable to the perturbation classes considered in our simulations, we summed elasticities over matrix elements corresponding to fecundity, adult survival, and juvenile survival, respectively. Fecundity elasticities were obtained by summing the elasticities of the non-zero elements of the fecundity matrix **F**. Survival elasticities were computed from the non-fecundity component of the projection matrix and partitioned according to whether the source stage corresponded to a reproductive or pre-reproductive class, yielding summed adult- and juvenile-survival elasticities. These aggregated elasticities were then related to the vulnerability metric Φ using univariate linear regressions across species.

## 3 Results

### 3.1 Heterogeneity in population trajectories across perturbation regimes

Species exhibited strongly heterogeneous demographic impacts across perturbation regimes, consistent with the idea that the effects of environmental forcing depend both on the vital rate under perturbation and on the underlying life-history strategy (Fig. 2). Representative trajectories under contrasting perturbation regimes spanned the full range from near-complete demographic resistance to rapid collapse, even when species were exposed to the same perturbation regime. In some cases, population size remained close to the unperturbed expectation throughout the simulation, whereas in others, identical perturbations produced pronounced declines from which recovery was only partial or entirely absent. Thus, the demographic consequences of environmental perturbations were highly context dependent, shaped by the interaction between perturbation attributes and species’ life-history traits.

**Figure 1:**
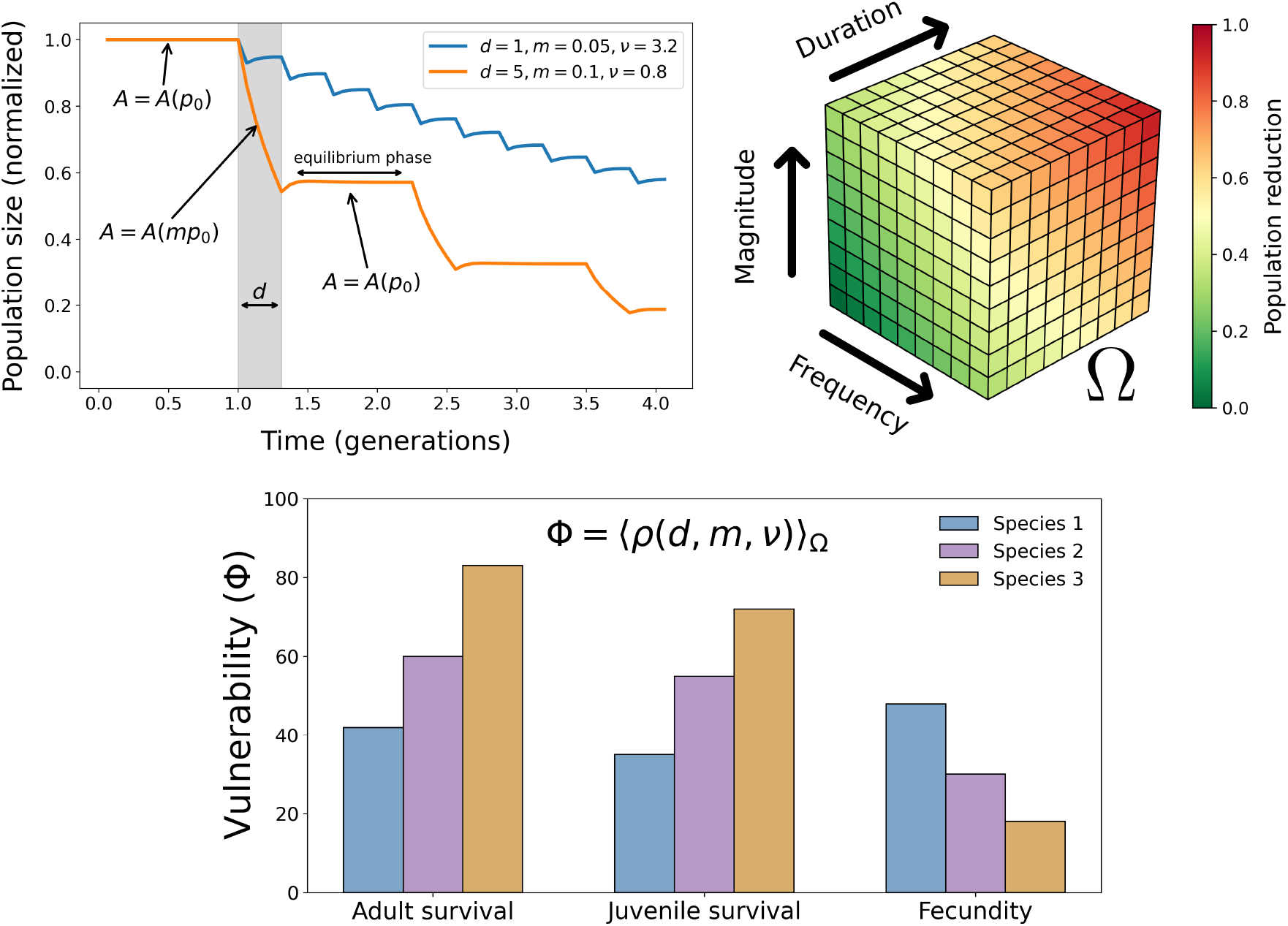
Schematic representation of the demographic vulnerability framework. We characterize perturbation regimes by three attributes: magnitude (*m*), representing the proportional reduction of a targeted demographic parameter; duration (*d*), with *d* = 1 corresponding to a pulse perturbation and *d >* 1 to a press perturbation; and frequency (*v*), defining the recurrence of perturbation events. For any combination of (*d, m, v*), population dynamics transition from the unperturbed state (*A* = *A*(*p*_0_)) to the perturbed state (*A* = *A*(*mp*_0_)), and, once the perturbation ceases, the system returns to the original projection matrix (*A* = *A*(*p*_0_)), potentially approaching a new equilibrium phase before the next event. Exploring the full perturbation space, {Ω}, defined by duration (*d*), magnitude (*m*), and frequency (*v*), allows us to compute the associated population reduction *ρ*(*d, m, v*), for each perturbation scenario. Vulnerability to perturbations affecting a given demographic parameter is then defined as the average population reduction over the full perturbation space, Φ = ⟨*ρ*(*d, m, v*)⟩_Ω_.

**Figure 2:**
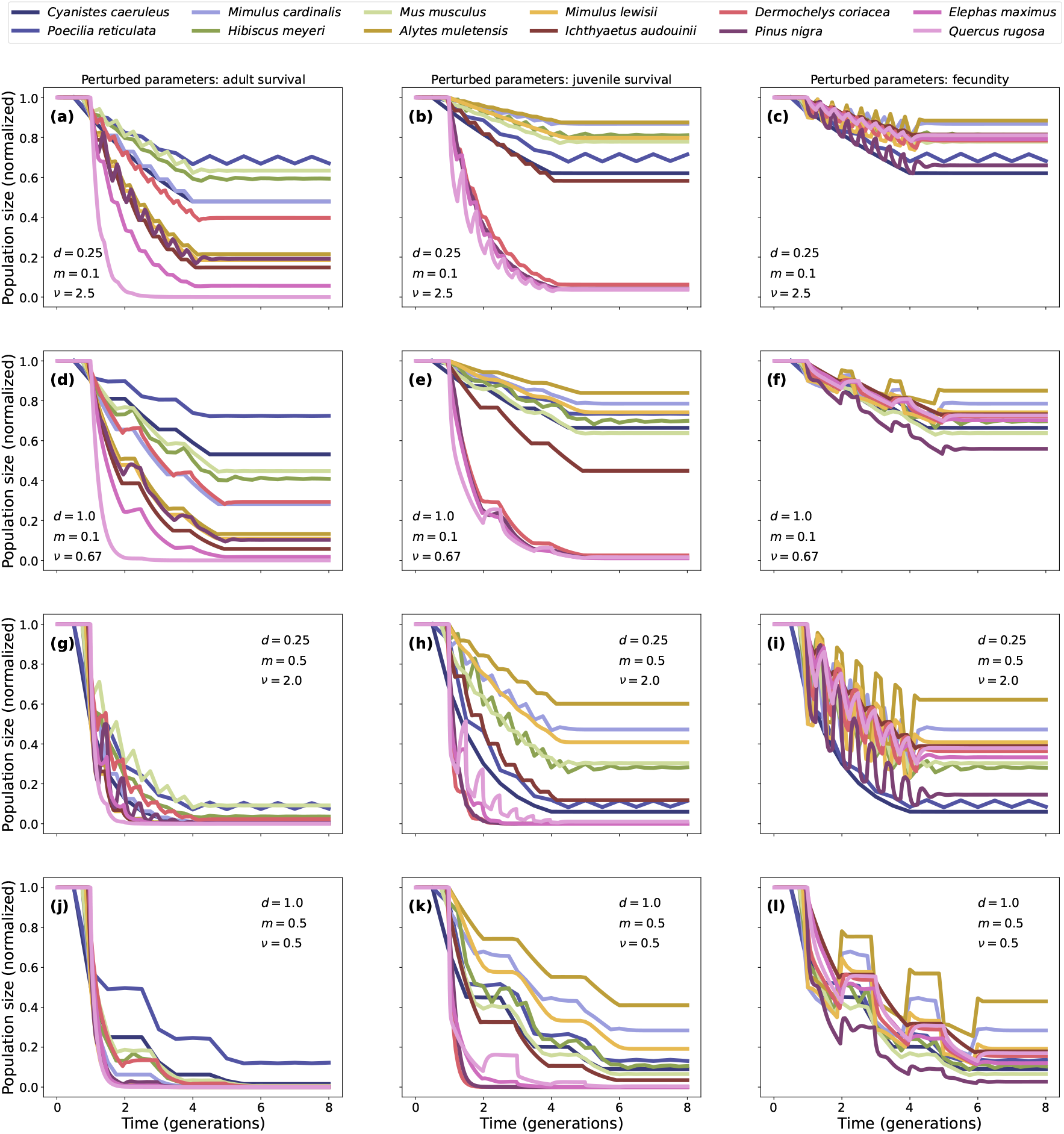
Population trajectories under contrasting perturbation regimes across species and demographic targets. Normalized population size is shown through time for the 12 study species under four representative perturbation regimes defined by magnitude (*m*), perturbation duration (*d*) and frequency (*v*). Columns correspond to perturbations affecting adult survival (a,d,g,j), juvenile survival (b,e,h,k), and fecundity (c,f,i,l). Rows represent four combinations of perturbation intensity and temporal structure: low-magnitude, short-duration, frequent perturbations (a–c; *m* = 0.1, *d* = 0.25, *v* = 2.5), low-magnitude, longer-duration perturbations, less frequent perturbations (d–f; *m* = 0.1, *d* = 1.0, *v* = 0.67), high-magnitude, short-duration perturbations, frequent perturbations (g–i; *d* = 0.25, *m* = 0.5, *v* = 2.0), and high-magnitude, longer-duration perturbations, less frequent perturbations (j–l; *d* = 1.0, *m* = 0.5, *v* = 0.5). The figure highlights the strong heterogeneity in demographic responses across species exposed to identical perturbation regimes. It also shows that sensitivity depends on the vital rate being perturbed: reductions in adult survival generally produce the strongest declines, whereas the relative ranking of species changes across demographic targets, illustrating the context dependence of vulnerability. Species are ordered in the legend based on generation time, 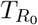, from left to right and from above to below.

In almost all species the strongest and most persistent impacts were generally observed when perturbations affected adult survival (Fig. 2a,d,g,j). Across a wide range of regimes, reductions in adult survival drove marked declines in population size and often pushed populations close to collapse. Perturbations to fecundity generally had weaker effects (Fig. 2c,f,i,l), with many species maintaining relatively large population sizes even under recurrent perturbations. In contrast, perturbations affecting juvenile survival generated a much broader spread of impacts across species (Fig. 2b,e,h,k), indicating that this life stage constitutes a major axis of demographic differentiation along the life-history continuum.

Beyond differences in final impact, the trajectories also revealed qualitatively distinct transient dynamics depending on the perturbed vital rate (Fig. 2). Perturbations affecting adult survival typically produced abrupt drops in population size followed by limited recovery between events, especially in slow-lived species, yielding rapid convergence towards strongly depleted states. By contrast, perturbations affecting juvenile survival often generated more delayed and cumulative declines, consistent with the progressive erosion of recruitment and the lagged propagation of juvenile losses through the life cycle. Fecundity perturbations, in turn, frequently produced oscillatory or saw-tooth dynamics synchronized with perturbation timing, but with comparatively modest long-term decline unless perturbations were both strong and recurrent. Thus, the trajectories differ not only in their final demographic impact, but also in their transient signatures, showing that different vital rates imprint distinct population dynamics.

### 3.2 Vulnerability across the perturbation space

To further understand the global impacts and population vulnerability to different species to perturbations, we explored the vulnerability metric across species. First to further illustrate how perturbation attributes interact with life-history strategies, we examined two-dimensional slices of the three-dimensional perturbation space obtained by fixing one perturbation attribute and varying the other two. As a first example, we studied the impact of a single-event perturbation of different magnitude and durations fixing frequency to *v* = 1*/*3, so that only one perturbation is applied over the whole simulation (Fig. 3). In this case, perturbations affecting adult survival generated the strongest demographic impacts across all representative strategies (Fig. 3a,d,g). Even moderate reductions in adult survival produced extensive regions of high population reduction, particularly in slow-lived species. In contrast to this pattern, perturbations affecting juvenile survival produced the greatest differentiation among life histories strategies (Fig. 3b,e,h), with fast-lived species remaining comparatively resistant and intermediate and slow-lived species showing much broader regions of high impact. Finally, perturbations affecting fecundity were, comparatively weak and more homogeneous (Fig. 3c,f,i).

**Figure 3:**
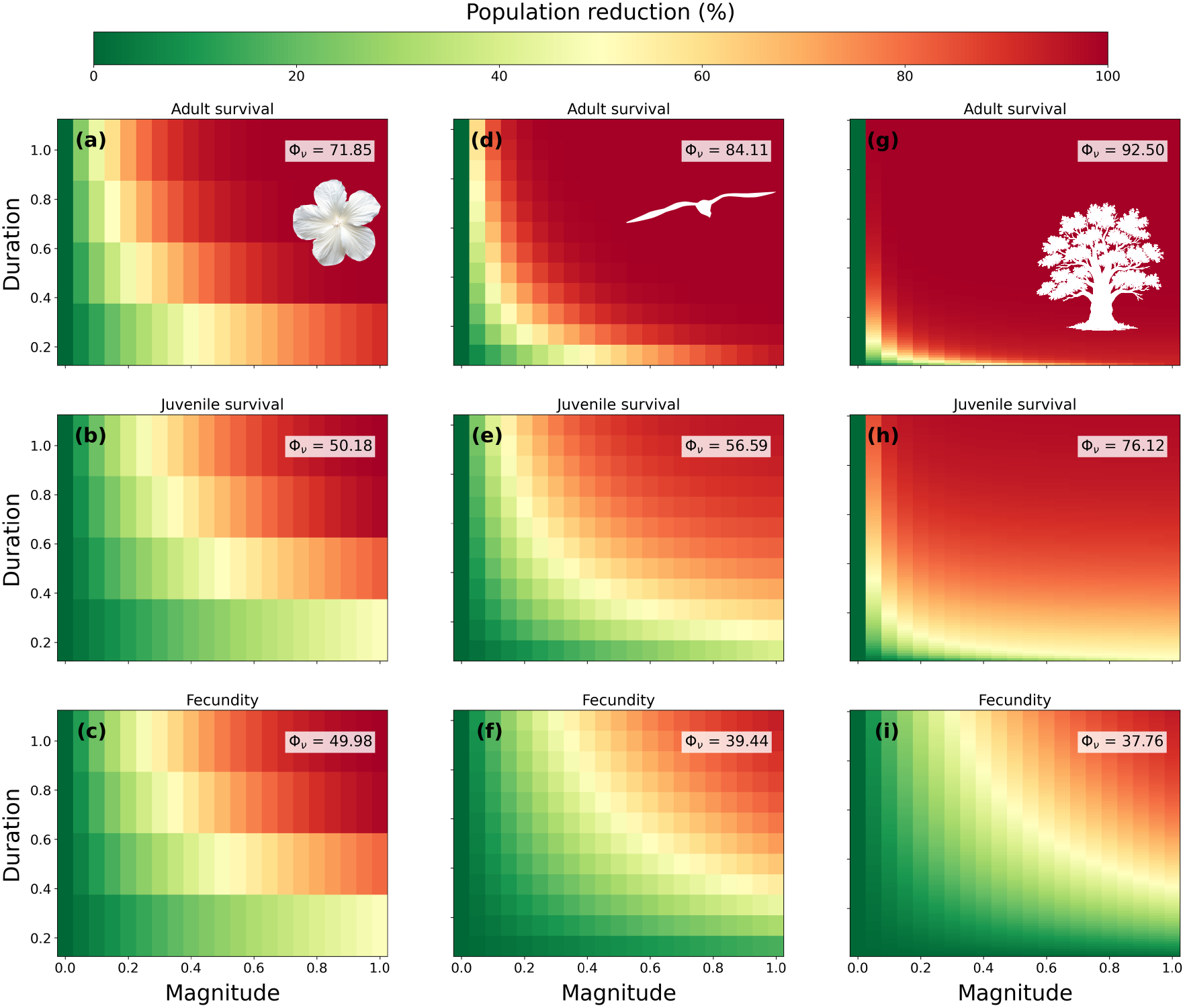
Population reduction across perturbation magnitude and duration for representative life histories at fixed frequency *v* = 1*/*3. Heatmaps show the percent population reduction, *ρ*(*m, d, v* = 1*/*3), for three species representing fast (*Hibiscus meyeri*, left column), intermediate (*Ichthyaetus audouinii*, middle column), and slow (*Quercus rugosa*, right column) life-history strategies. Perturbations were applied at fixed frequency to adult survival (top row), juvenile survival (middle row), and fecundity (bottom row). Colours range from low demographic impact (green) to near-complete collapse (red). Values of Φ_*v*_ in each panel indicate the mean population reduction across the displayed (*m, d*) slice at fixed *v* = 1*/*3. The number of bins in the duration space (i.e., the resolution of the Y axis) are constrained by the generation time of the species, as the maximum number of time-steps corresponding to *d* = 1 is the generation time of the species.

Similar conclusions emerged when examining regimes with fixed perturbation magnitude at *m* = 0.1, this is, 10% reduction in the the perturbed vital rate. Impacts increased sharply with perturbation duration and recurrence, with perturbations affecting adult survival generating the highest impact, particularly in slow-lived species. Again, impacts on juvenile survival show the highest degree od differentiation among the species, while impacts to fecundity were more homogeneous (Supplementary Fig. 1). When perturbation duration was held constant at *d* = 1, similar results were obtained regarding adult and juvenile survival, but a change of tendency emerged for fecundity, where fast-lived species became more vulnerable than slow-lived ones Supplementary Fig. 2).

These insights are representative of just a specific subset of the full perturbation space in which a given perturbation dimension (duration, magnitude or frequency) is kept constant. Then, we analyze the overall vulnerability index to summarize how the different species are affected by the full perturbation space through the integrated vulnerability metric. The overall vulnerability preserves the same qualitative ordering previously observed (Fig. 4): vulnerability is highest for perturbations affecting adult survival, most variable for perturbations affecting juvenile survival, and lowest overall for perturbations affecting fecundity.

**Figure 4:**
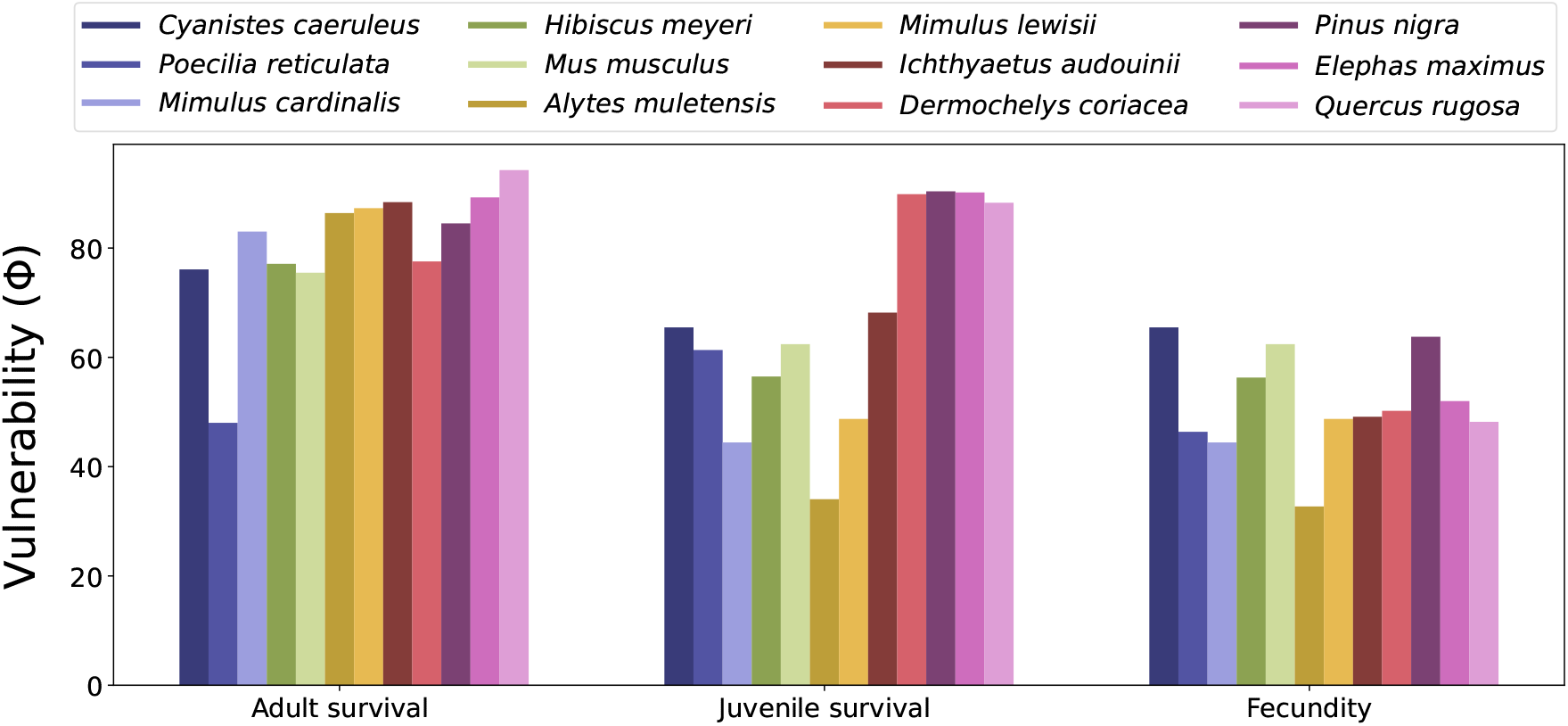
Integrated vulnerability across perturbed demographic parameters. Bars show the vulnerability metric, Φ, for perturbations affecting adult survival, juvenile survival, and fecundity. Higher values indicate greater overall demographic vulnerability after integrating over all simulated perturbation regimes. Species are ordered in the legend and in the x axis based on generation time, 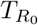, being *Cyanistes caeruleus* and *Poecilia reticulata* those with shorter generation times and *Elephas maximus* and *Quercus rugosa* those with the larger ones

Together, all these complementary perspectives converge on a consistent picture: adult survival is the dominant axis of demographic vulnerability, juvenile survival is the dominant axis of differentiation among life-history strategies, and fecundity is comparatively buffered.

### 3.3 Distinct life-history descriptors predict vulnerability to survival and reproductive perturbations

Generation time, 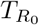, captured a substantial fraction of the among-species variation in integrated vulnerability, but only when perturbations acted on survival. Vulnerability to perturbations affecting adult and juvenile survival increased significantly with generation time (*R*^2^ = 0.42, *p* = 2.25 ×10^*−*2^, and *R*^2^ = 0.43, *p* = 2.05 × 10^*−*2^, respectively; Fig. 5a,b), indicating that slow life histories are consistently more affected to survival reduction. By contrast, vulnerability to fecundity perturbations was unrelated to generation time (*R*^2^ = 0.02, *p* = 0.65; Fig. 5c), suggesting that sensitivity to reproductive failure does not follow the fast–slow continuum.

**Figure 5:**
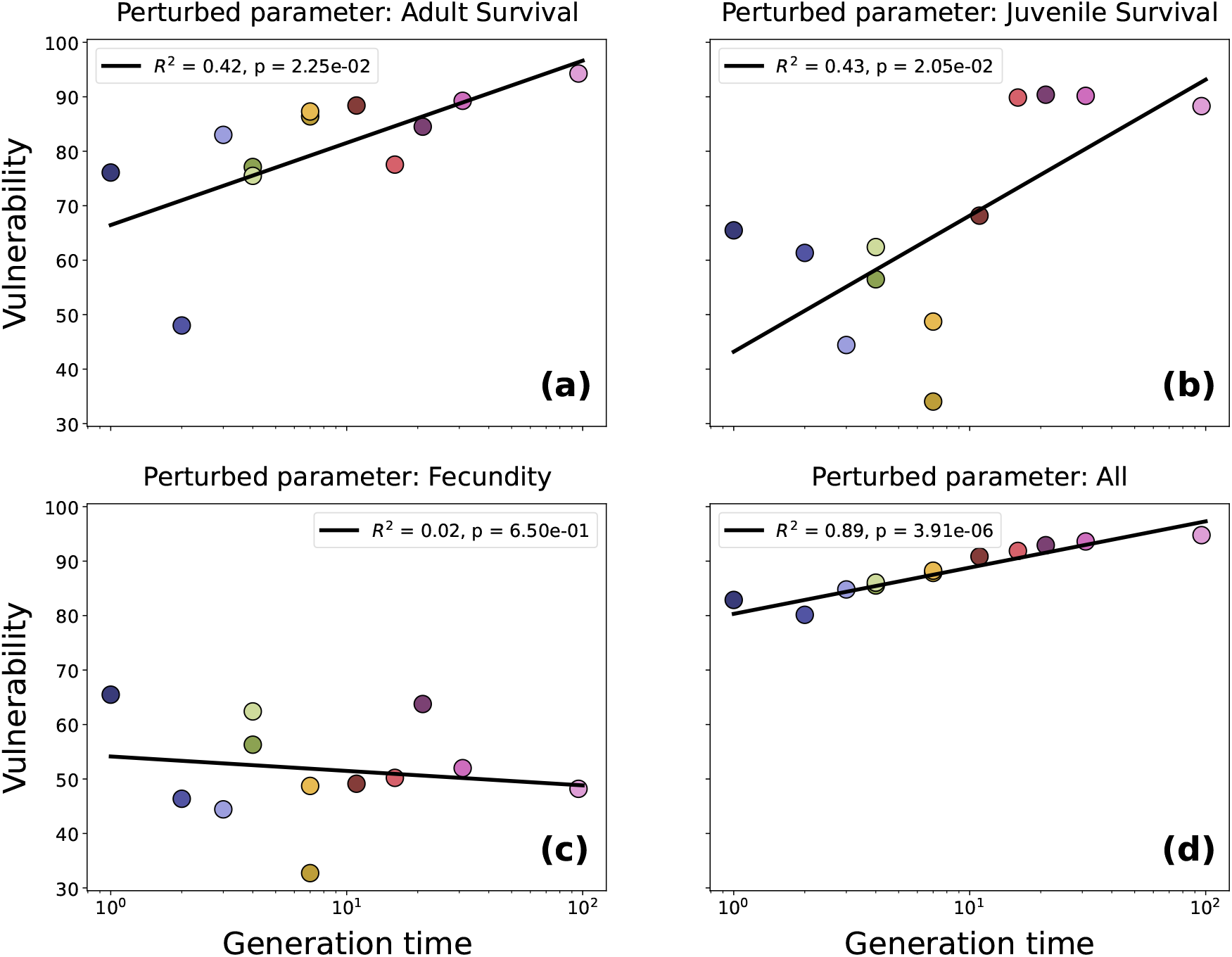
Relationship between population vulnerability and species generation time. Scatter plots show the relationship between vulnerability (Φ) and generation time 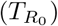 across species for perturbations affecting (a) adult survival, (b) juvenile survival, and (c) fecundity. Panel (d) shows vulnerability to perturbations applied to all parameters. Lines indicate linear regressions, with *R*^2^ and *p*-values showing the strength and significance of the relationship. Vulnerability increases with generation time when perturbations affect survival, but not when they affect fecundity. When all perturbation types are combined, generation time becomes a strong predictor of overall demographic vulnerability. Species are represented by the same colors as in Fig. 3

When vulnerability was aggregated across all perturbed demographic parameters, the relationship with generation time became stronger (*R*^2^ = 0.89, *p* = 3.91 × 10^*−*6^; Fig. 5d). Thus, although the effect of life-history pace depends on which vital rate is perturbed, generation time emerges as a powerful predictor of overall demographic vulnerability. Long-lived species are not inherently more vulnerable to every type of perturbation, yet the perturbations that generate the largest demographic impacts disproportionately strike the vital rates most critical to their persistence. Consequently, even if vulnerability is not uniformly higher in the slow end of the continuum, these species are disproportionately affected by the perturbation regimes that dominate the integrated demographic impact.

The demographic traits associated with vulnerability differed sharply among perturbed vital rates, indicating that survival and fecundity perturbations are filtered through different dimensions of life-history variation. For perturbations affecting adult survival, vulnerability was most strongly associated with descriptors capturing the temporal organization of the life cycle, with entropy and generation-time metrics emerging as the strongest predictors (Fig. 6a). Vulnerability to perturbations affecting juvenile survival was explained by a broader set of temporal descriptors, including age at reproduction, life expectancy, generation time, and entropy-related measures (Fig. 6b). Thus, vulnerability to survival perturbations is governed primarily by how demographic events are distributed across the life cycle and by how quickly populations can replace lost individuals. A different picture emerged for fecundity perturbations. Here, vulnerability was best predicted by reproductive output, with *R*_0_ and 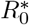 clearly outperforming the remaining descriptors (Fig. 6c). This result is consistent with the weak relationship between fecundity vulnerability and generation time: resistance to reproductive failure depends less on life-history pace than on baseline reproductive capacity. In other words, species differ little in their sensitivity to fecundity perturbations along the fast–slow continuum, but differ more in how much reproductive output they can lose before population reduction becomes substantial.

**Figure 6:**
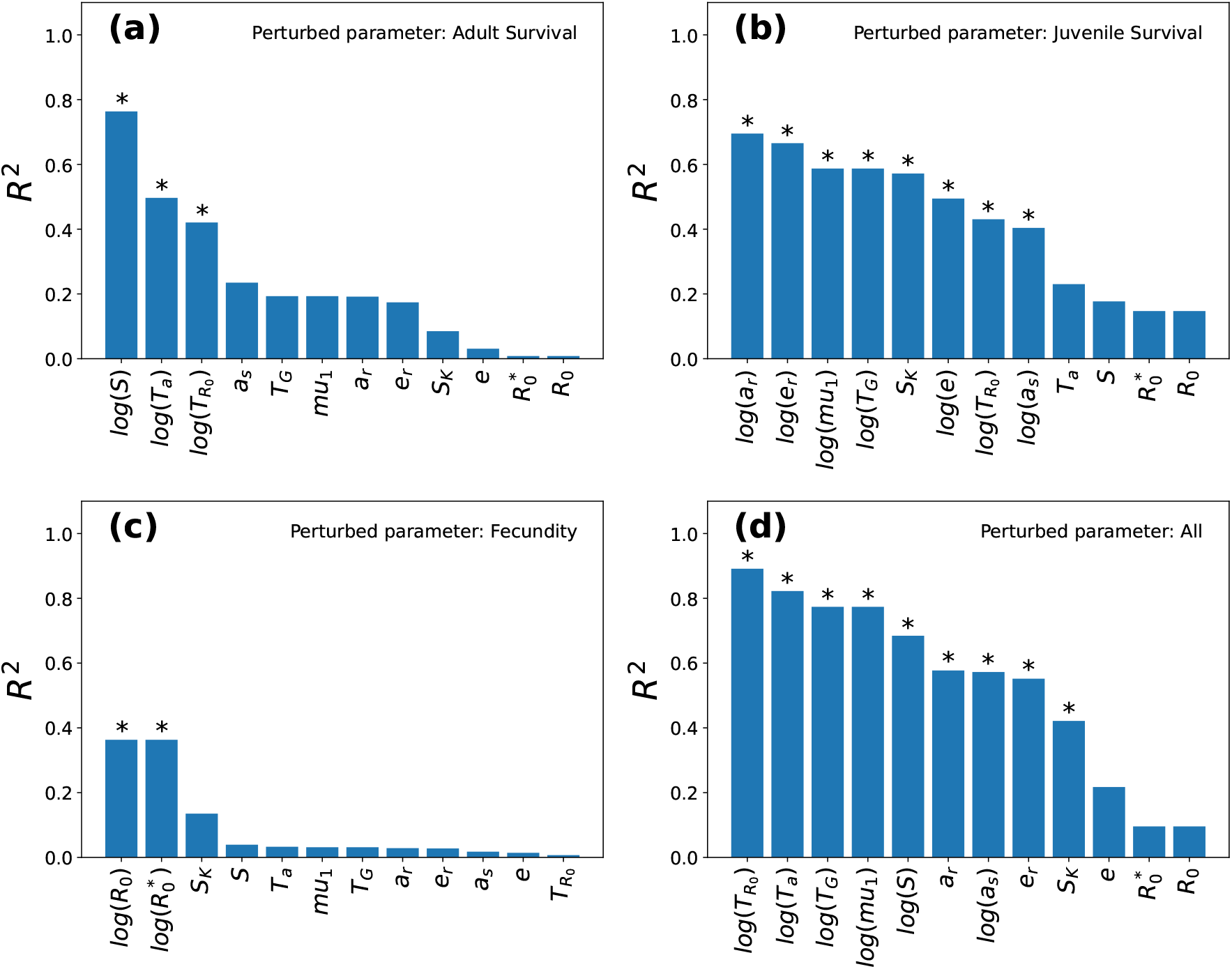
Life-history predictors of integrated vulnerability depend on the perturbed vital rate. Bars show the explanatory power (*R*^2^) of demographic descriptors for the vulnerability metric (Φ) across perturbations affecting adult survival (a), juvenile survival (b), fecundity (c), and all demographic parameters combined (d). Descriptors include reproductive output 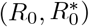, generation-time metrics 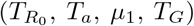, life expectancy (*e, e*_*r*_), age structure (*a*_*s*_), age at reproduction (*a*_*r*_), and entropy measures (*S, S*_*K*_). Asterisks denote statistically significant associations (*p <* 0.05). Survival-related vulnerability is best predicted by temporal and entropy-related descriptors, whereas fecundity vulnerability is primarily predicted by reproductive output.

When all demographic parameters were perturbed at the same time, generation-time and entropy-related descriptors again ranked highest (Fig. 6d). This indicates that, despite the distinct demographic filter associated with fecundity perturbations, overall vulnerability across the perturbation space is ultimately dominated by the temporal structure of the life cycle. Taken together, these results show that the demographic determinants of vulnerability are parameter specific, but that slow, temporally concentrated life histories remain the most exposed to the cumulative effects of environmental perturbation.

The strongest relationships make this contrast explicit (Supplementary Fig. 3): entropy best predicted vulnerability to adult-survival perturbations, age at reproduction best predicted vulnerability to juvenile-survival perturbations, reproductive output best predicted vulnerability to fecundity perturbations, and generation time best predicted aggregate vulnerability. Thus, no single life-history axis captures all forms of demographic vulnerability.

The elasticity of the perturbed demographic parameters was positively associated with vulnerability, but explained only part of its variation (Supplementary Fig. 4). Elasticity accounted for 52% and 58% of the variance in vulnerability for adult- and juvenile-survival perturbations, respectively, and only 20% for fecundity. Thus, although Φ is related to classical asymptotic perturbation metrics, it also captures additional information, most likely because it integrates the effects of perturbation magnitude, duration, and frequency, and therefore the temporal structure of perturbation regimes. Consistent with this, several life-history descriptors explained more variation in vulnerability than elasticity alone.

## 4 Discussion

Our study introduces a unified demographic framework to quantify the vulnerability of biological populations to a wide spectrum of environmental perturbations. By decomposing disturbances into magnitude, duration, and frequency, we capture the full continuum of perturbation regimes, rather than treating pulse and press disturbances as separate categories. In doing so, the framework moves beyond traditional sensitivity and elasticity analyses, which quantify the effects of small changes in vital rates on asymptotic population growth, by explicitly capturing how perturbations of different magnitude, duration, and frequency interact with life-history strategy to shape demographic vulnerability. This provides a more complete mechanistic perspective on how life-history strategies shape population persistence under variable environmental pressures. Building on this framework, we introduce a vulnerability metric, Φ, as a systematic way to quantify population persistence to different perturbation regimes into a single measure. This parsimonious approach synthesizes complex demographic impacts into a comparable value and highlights general principles linking environmental variability with the evolution of life-history strategies [14].

A primary finding of this work is that all analyzed species are highly vulnerable to perturbations in adult survival: across the fast-slow continuum, reductions in adult survival led to the most severe population declines. Aligned with established demographic theory, which posits that adult survival often has the highest elasticity in long-lived species [46], our framework also confirms that long-lived species are the most vulnerable to perturbations in adult survival. However, even for “fast” species, sustained adult mortality remains a critical threat that can quickly overwhelm reproductive compensation. Conversely, all the species are much less vulnerable to perturbations in fecundity. In contrast, juvenile survival emerged as the key demographic parameter for differentiating life-history strategies. The high heterogeneity of perturbations to juvenile survival indicates that this life stage may constitute a major axis of demographic variation across life-history strategies [47]. Slower-lived species, which often exhibit lower reproductive rates and later maturation, proved to be more impacted by juvenile losses, leading to the maximal spread in the vulnerability metric Φ observed in our results. The comparatively greater homogeneity in vulnerability to fecundity perturbations relative to juvenile perturbations, suggests that reproductive perturbations or failures may be a relatively common, and that most species are comparatively resilient to fluctuations in reproductive output.

The quantification of vulnerability through the Φ metric offers valuable insights for evolutionary ecology. By condensing complex demographic impacts into a single, comparable measure, we provide a unified way to evaluate how different species are affected across the full spectrum of perturbation regimes. From an evolutionary perspective, it helps clarify the selection pressures that may have shaped life-history strategies under different disturbance environments. In environments with frequent or unpredictable shocks to survival, fast strategies are expected to be favored because they allow rapid demographic replacement. By contrast, slow strategies might only evolve in ecological contexts where adult survival is maintained consistently high, either because the environment naturally imposes few disturbances on survival or because the conditions allow the gradual evolution of traits that buffer adult survival [48–50]. More generally, high vulnerability of a given life-history strategy to perturbations affecting a particular demographic process may indicate either that such perturbations have historically been rare in the environments where that strategy evolved, or that selection has favored buffering mechanisms that reduce variation in the affected demographic parameter.

An important feature of our framework is that perturbation regimes are defined relative to generation time rather than absolute time [4, 13]. This allows species with very different life histories to be compared over equivalent demographic windows, but it also means that the same perturbation duration can correspond to very different absolute times for fast- and slow-lived species. Our results should therefore be interpreted primarily as a comparative description of vulnerability relative to life-history pace, rather than as a direct assessment of risk over fixed management horizons. Even so, the approach may guide conservation by identifying species with higher vulnerability to perturbations globally or affecting particular demographic parameters.

Additionally, the framework offers substantial potential for other conservation applications. By restricting analyses to specific regions of the perturbation space, focusing on ecologically relevant disturbance regimes, or redefining perturbation duration in calendar rather than generational time, the metric could provide a practical tool for evaluating the demographic impacts of particular perturbation scenarios. Such applications could facilitate comparisons of demographic vulnerability across species exposed to similar disturbance regimes and help assess how changes in disturbance intensity, duration, or recurrence under alternative management scenarios may influence projected population responses [37].

Our framework complements several recent lines of work on ecological perturbation and demographic response. Studies of transient dynamics and demographic resilience have also shown that short-term responses to disturbance depend strongly on life-history structure and that can be decomposed into components such as resistance, compensation, and recovery [24, 25, 39]. Related work has also shown that pulse disturbances in age-structured populations generate life-history-dependent impacts and recovery times [26], and that pulse and press disturbances can differ markedly in their transient consequences even when long-term outcomes are similar [27]. More recently, work on perturbation variability and structured demographic buffering has emphasized that temporal environmental structure itself can shape ecological and demographic responses [28, 29]. Our contribution differs from these approaches by integrating perturbation magnitude, duration, and frequency within a single demographic space and by comparing how perturbations acting on different vital rates are filtered across the fast–slow continuum.

While our framework provides a clear comparative baseline, its intentional simplicity also imposes several limitations. For instance, we do not include density-dependent feedbacks or compensatory demographic responses. In natural populations, declines in one vital rate can be partly offset by favorable changes in others, for example, reduced adult density may relax competition and enhance juvenile survival or fecundity, thereby buffering population decline [51]. Similarly, physiological constraints, carry-over effects, and life-history trade-offs can create delayed or state-dependent coupling among vital rates, so that perturbations affecting one demographic process may propagate to others across subsequent time steps. Capturing these effects will require models in which vital rates depend explicitly on one another, or on internal state variables, rather than being perturbed independently [11, 52]. Additionally, except when all vital rates are perturbed simultaneously, we considered vital rates as independent targets of perturbation, whereas environmental forcing may affect multiple demographic processes at the same time [4, 53]. Such within-time-step covariation among vital rates can modify population dynamics and should ideally be incorporated through covariance structures linking demographic parameters (e.g. survival and reproduction) [54]. Moreover, because we perturb vital rates directly rather than the environmental drivers that shape them, our results isolate intrinsic demographic sensitivity rather than realized responses to specific environmental scenarios. Some of these limitations are common to current demographic resilience frameworks [25], and the main contribution of our approach is not to reproduce the full complexity of real population dynamics, but rather to isolate the core demographic dimensions through which vulnerability is structured across life histories.

Although we attempted to capture a broad range of generation times, life-history strategies, and taxonomic groups spanning both plants and vertebrates, important groups such as invertebrates remain absent from our analysis. Applying the framework to encompass this broader diversity of life histories represents an important avenue for future work. An important challenge for future work will be to incorporate life-history trade-offs and demographic compensation and covariance into this framework. Extending vulnerability metrics in this direction would allow to capture not only the direct demographic impacts of perturbations, but also the broader capacity of species to respond to disturbance across life-history strategies and perturbation regimes. An additional challenge will be to embed this targeted perturbation framework within environmentally driven stochastic demographic models, allowing perturbation strength to be evaluated relative to the natural variability experienced by each species [3–6]. Finally, scaling up the framework to ecological communities would make it possible to study how temporally structured perturbations propagate through multispecies systems, thereby linking our demographic approach to recent nonequilibrium theory on community responses to pulse perturbations [55].

Overall, our study provides a basic framework for understanding the rules of life that govern how biological diversity persists in an ever-changing world. This framework is particularly relevant under ongoing global change, which increasingly exposes species to environmental conditions outside the range under which they evolved. By revealing how species with contrasting life-history strategies differ in their capacity to withstand such novel perturbations, our work offers insights into fundamental principles of demographic vulnerability.

## Data and code accessibility

All matrix population models used in this study, including the corresponding CSV files employed in the analyses, are available as supplementary material. To support the assessment of demographic vulnerability in structured populations, we developed open-source implementations of the vulnerability framework introduced here in R [56], Python [57], and Julia [58]. The R package `demovuln-r` is available through [GitHub](https://github.com/agimenezromero/demovuln-r) and has been submitted to CRAN; the Python package `demovuln` is available through [GitHub](https://github.com/agimenezromero/demovuln), [PyPI](https://pypi.org/project/demovuln/), and online [documentation](https://demovuln.readthedocs.io/en/latest/); and the Julia package `DemoVuln.jl` is available through [GitHub](https://github.com/agimenezromero/DemoVuln.jl).

## Acknowledgements

AGR acknowledges financial support from grant JDC2024-053275-I, funded by MICIU/AEI/10.13039/501100011033 and FSE+. AGR and MG acknowledge financial support from the Spanish Ministerio de Ciencia e Innovación/AEI and EU-FEDER (PID2021-124731NB-I00 and PIE 202430E234).

