## Supplementary material for "A demographic framework for assessing population vulnerability to contrasting perturbation regimes": SI.pdf

### 1 Matrix Population Models

This section provides the matrix population models (MPMs) employed in the analyses presented in the main text. For each species, we report the projection matrix **A** used in the simulations, with rows representing the production of individuals in each stage at time $t+1$ and columns representing the contribution of each stage at time $t$. These matrices were used both to compute the life-history descriptors and to define the baseline demographic dynamics before perturbations were imposed.

Because the dimensionality of the MPMs differs markedly among species, the matrices are shown using a compact notation. Small matrices are displayed directly, whereas for larger age- or stage-structured matrices we provide only the reference. In any case, we provide CSV files for each of the matrices in the Zenodo repository.

#### 1.1 *Alytes muletensis*

The projection matrix used for *Alytes muletensis* was a $3 \times 3$ stage-structured matrix was obtained from the COMADRE database [1, 2],

$$\mathbf{A} = \begin{pmatrix} 0 & 0 & 6.86 \\ 0.003 & 0 & 0 \\ 0.007 & 0.18 & 0.73 \end{pmatrix} \tag{1}$$

This matrix was used as the unperturbed demographic operator for all simulations involving this species.

#### 1.2 *Cyanistes caeruleus*

The projection matrix used for *Cyanistes caeruleus* was a $5 \times 5$ stage-structured matrix obtained from the COMADRE database [3],

$$\mathbf{A} = \begin{pmatrix} 0.65 & 0.69 & 0.69 & 0.59 & 0.59 \\ 0.34 & 0 & 0 & 0 & 0 \\ 0 & 0.34 & 0 & 0 & 0 \\ 0 & 0 & 0.34 & 0 & 0 \\ 0 & 0 & 0 & 0.34 & 0.34 \end{pmatrix} \tag{2}$$

This matrix was used as the unperturbed demographic operator for all simulations involving this species.

##### 1.3 *Dermochelys coriacea*

The projection matrix used for *Dermochelys coriacea* was a $8 \times 8$ stage-structured matrix obtained from [4], given by

$$\mathbf{A} = \begin{pmatrix} 0 & 0 & 0 & 10.7969 & 0 & 0 & 10.7969 & 0 \\ 0.465 & 0.3174 & 0 & 0 & 0 & 0 & 0 & 0 \\ 0 & 0.1476 & 0.8325 & 0 & 0 & 0 & 0 & 0 \\ 0 & 0 & 0.0526 & 0.0041 & 0.2428 & 0.4825 & 0.6083 & 0 \\ 0 & 0 & 0 & 0.8889 & 0 & 0 & 0 & 0 \\ 0 & 0 & 0 & 0 & 0.6502 & 0 & 0 & 0 \\ 0 & 0 & 0 & 0 & 0 & 0.4105 & 0.2847 & 0 \\ 0 & 0 & 0.0079 & 0 & 0 & 0 & 0 & 0 \end{pmatrix} \tag{3}$$

This matrix was used as the unperturbed demographic operator for all simulations involving this species.

##### 1.4 *Elephas maximus*

The projection matrix used for *Elephas maximus* is a $60 \times 60$ stage-structured matrix obtained from the COMADRE database [5, 6].

##### 1.5 *Hibiscus meyeri*

The projection matrix used for *Hibiscus meyeri* was a $26 \times 26$ stage-structured matrix obtained from [7].

##### 1.6 *Ichthyaetus audouinii*

The projection matrix used for *Ichthyaetus audouinii* was a $7 \times 7$ stage-structured matrix obtained from [8, 9], given by

$$\mathbf{A} = \begin{pmatrix} 0 & 0 & 0.0975 & 0 & 0 & 0 & 0.1748 \\ 0.744 & 0 & 0 & 0 & 0 & 0 & 0 \\ 0 & 0.4579 & 0 & 0 & 0 & 0 & 0 \\ 0 & 0.4881 & 0 & 0 & 0 & 0 & 0 \\ 0 & 0 & 0 & 0.2948 & 0 & 0 & 0 \\ 0 & 0 & 0 & 0 & 0.5996 & 0 & 0 \\ 0 & 0 & 0.924 & 0.6292 & 0.2914 & 0.891 & 0.891 \end{pmatrix} \tag{4}$$

This matrix was used as the unperturbed demographic operator for all simulations involving this species.

##### 1.7 *Mimulus cardinalis*

The projection matrix used for *Mimulus cardinalis* was a $4 \times 4$ stage-structured matrix obtained from [10],

$$\mathbf{A} = \begin{pmatrix} 0.199 & 802 & 5.82e+03 & 3.05e+04 \\ 2.66 \times 10^{-5} & 0.0776 & 0.0231 & 0.0011 \\ 7.94 \times 10^{-6} & 0.0807 & 0.322 & 0.216 \\ 2.91 \times 10^{-7} & 0.0158 & 0.115 & 0.601 \end{pmatrix} \tag{5}$$

This matrix was used as the unperturbed demographic operator for all simulations involving this species.

##### 1.8 *Mimulus lewisii*

The projection matrix used for *Mimulus lewisii* was a $4 \times 4$ stage-structured matrix obtained from [10],

$$\mathbf{A} = \begin{pmatrix} 0.0325 & 222.5 & 8.61e+03 & 4.84e+04 \\ 2.01 \times 10^{-5} & 0.0827 & 0.0197 & 0 \\ 8.39 \times 10^{-6} & 0.0896 & 0.5451 & 0.1424 \\ 1.1 \times 10^{-7} & 0.0039 & 0.1527 & 0.7534 \end{pmatrix} \tag{6}$$

This matrix was used as the unperturbed demographic operator for all simulations involving this species.

##### 1.9 *Mus musculus*

The projection matrix used for *Mus musculus* was a $15 \times 15$ stage-structured matrix obtained from the COMADRE database [11].

##### 1.10 *Pinus nigra*

The projection matrix used for *Pinus nigra* was a $8 \times 8$ stage-structured matrix obtained from the COMPADRE database [12, 13],

$$\mathbf{A} = \begin{pmatrix} 0 & 0 & 0 & 0 & 0 & 0 & 145 & 821.28 \\ 0.63 & 0 & 0 & 0 & 0 & 0 & 0 & 0 \\ 0 & 0.63 & 0 & 0 & 0 & 0 & 0 & 0 \\ 0 & 0 & 0.63 & 0 & 0 & 0 & 0 & 0 \\ 0 & 0 & 0 & 0.63 & 0 & 0 & 0 & 0 \\ 0 & 0 & 0 & 0 & 0.63 & 0.915 & 0 & 0 \\ 0 & 0 & 0 & 0 & 0 & 0.069 & 0.915 & 0 \\ 0 & 0 & 0 & 0 & 0 & 0 & 0.069 & 0.983 \end{pmatrix} \tag{7}$$

This matrix was used as the unperturbed demographic operator for all simulations involving this species.

##### 1.11 *Poecilia reticulata*

The projection matrix used for *Poecilia reticulata* was a $14 \times 14$ stage-structured matrix obtained from the COMADRE database [14].

##### 1.12 *Quercus rugosa*

The projection matrix used for *Quercus rugosa* was a $7 \times 7$ stage-structured matrix obtained from the COMPADRE database [15],

$$
\mathbf{A} = \begin{pmatrix} 0 & 0 & 0 & 0 & 0.21 & 0.32 & 0.295 \\ 0.613 & 0.879 & 0.08 & 0 & 0 & 0 & 0 \\ 0 & 0.061 & 0.784 & 0.097 & 0 & 0 & 0 \\ 0 & 0 & 0.094 & 0.808 & 0 & 0 & 0 \\ 0 & 0 & 0 & 0.052 & 0.951 & 0 & 0 \\ 0 & 0 & 0 & 0 & 0.0306 & 0.9491 & 0 \\ 0 & 0 & 0 & 0 & 0 & 0.0447 & 0.985 \end{pmatrix} \tag{8}
$$

This matrix was used as the unperturbed demographic operator for all simulations involving this species.

### 2 Life history descriptors

| | *Cyanistes caeruleus* | *Poecilia reticulata* | *Mimulus cardinalis* | *Hibiscus meyeri* | *Mus musculus* | *Alytes muletensis* | *Mimulus lewisii* | *Ichthyaetus audouinii* | *Dermochelys coriacea* | *Pinus nigra* | *Elephas maximus* | *Quercus rugosa* |
|---|---|---|---|---|---|---|---|---|---|---|---|---|
| $R_0$ | 1.00 | 43.99 | 0.53 | 18.25 | 31.91 | 0.19 | 1.18 | 1.05 | 1.52 | 3296.84 | 5.17 | 1.17 |
| $R_0^*$ | 1.00 | 43.99 | 0.53 | 18.25 | 31.91 | 0.19 | 1.18 | 1.05 | 1.52 | 3296.84 | 5.17 | 1.17 |
| $T_{R_0}$ | 1.52 | 2.46 | 3.95 | 4.17 | 4.22 | 7.25 | 7.50 | 11.97 | 16.26 | 21.83 | 31.92 | 96.58 |
| $T_a$ | 1.52 | 2.05 | 5.02 | 3.46 | 2.78 | 13.12 | 7.19 | 11.83 | 15.16 | 11.84 | 28.20 | 93.11 |
| $T_G$ | 1.52 | 4.57 | 3.18 | 5.85 | 7.24 | 4.78 | 7.83 | 12.11 | 17.49 | 84.90 | 36.48 | 100.28 |
| $\mu_1$ | 1.52 | 4.57 | 3.18 | 5.85 | 7.24 | 4.78 | 7.83 | 12.11 | 17.49 | 84.90 | 36.48 | 100.28 |
| $a_r$ | 1.37 | 4.36 | 1.00 | 4.31 | 7.01 | 3.43 | 1.00 | 9.56 | 13.30 | 50.61 | 33.49 | 65.61 |
| $e$ | 1.52 | 9.73 | 1.00 | 2.86 | 12.00 | 1.03 | 1.00 | 8.41 | 2.58 | 8.40 | 35.00 | 15.50 |
| $e_r$ | 1.73 | 9.82 | 1.00 | 5.83 | 12.02 | 4.78 | 1.00 | 15.09 | 17.78 | 76.28 | 51.45 | 100.70 |
| $a_s$ | 1.52 | 1.27 | 1.00 | 1.43 | 1.78 | 2.26 | 1.00 | 8.90 | 4.06 | 1.86 | 14.36 | 34.01 |
| $S$ | 1.41 | 0.26 | 6.89 | 3.65 | 1.76 | 5.40 | 8.19 | 6.08 | 12.28 | 7.43 | 5.08 | 34.17 |
| $S_K$ | 0.56 | 0.23 | 0.00 | 1.09 | 0.14 | 0.17 | 0.00 | 1.03 | 1.38 | 2.49 | 0.64 | 1.79 |

#### 3 Results

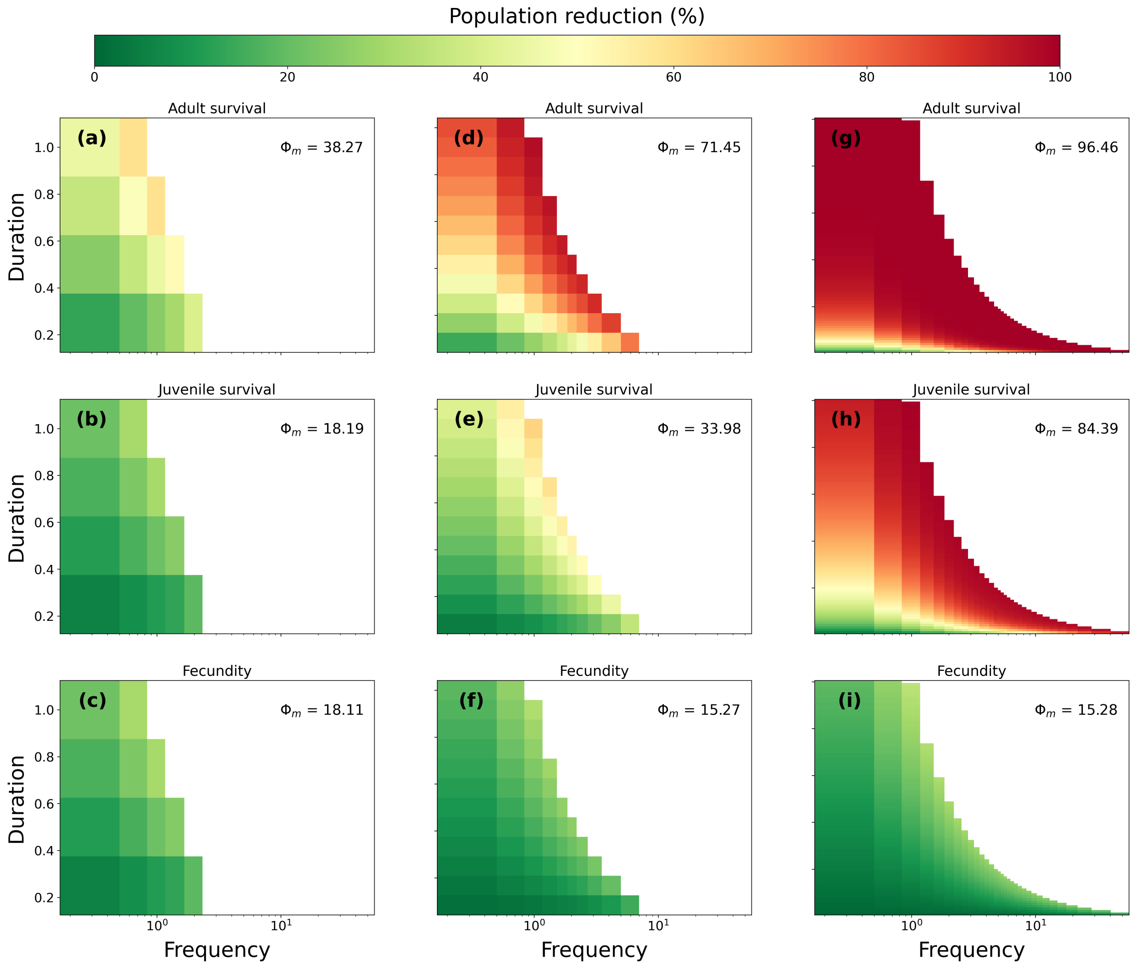


**Figure 1: Population reduction across perturbation duration and frequency at fixed magnitude.** Heatmaps show the percent population reduction, $\rho(m, d, \nu)$, for three species representing fast (left column), intermediate (middle column), and slow (right column) life-history strategies, with perturbation magnitude held constant at $m = 0.1$. Rows correspond to perturbations affecting adult survival, juvenile survival, and fecundity. Colors range from low demographic impact (green) to near-complete collapse (red). Values in each panel indicate the mean population reduction across the displayed $(d, \nu)$ slice. Blank regions indicate infeasible perturbation regimes, where perturbation duration exceeds the interval between successive events or perturbations would occur more than once per time step.

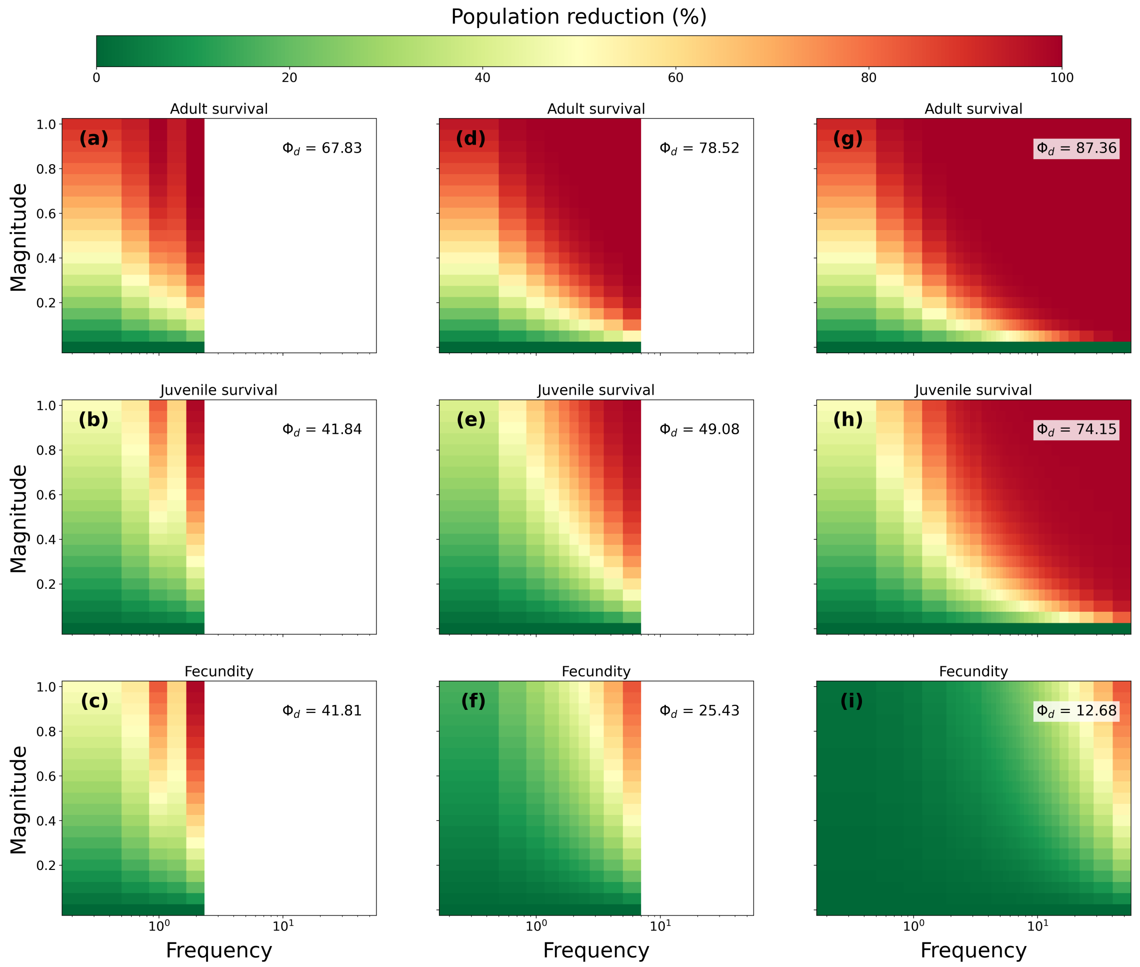


**Figure 2: Population reduction across perturbation magnitude and frequency at fixed duration.** Heatmaps show the percent population reduction, $\rho(m, d, \nu)$, for three species representing fast (left column), intermediate (middle column), and slow (right column) life-history strategies, with perturbation duration held constant at $d = 1$. Rows correspond to perturbations affecting adult survival, juvenile survival, and fecundity. Colors range from low demographic impact (green) to near-complete collapse (red). Values in each panel indicate the mean population reduction across the displayed $(m, \nu)$ slice. These slices recover the same qualitative ordering observed in the main text, while also showing that recurrent, high-magnitude fecundity perturbations can be especially costly in fast-lived species.

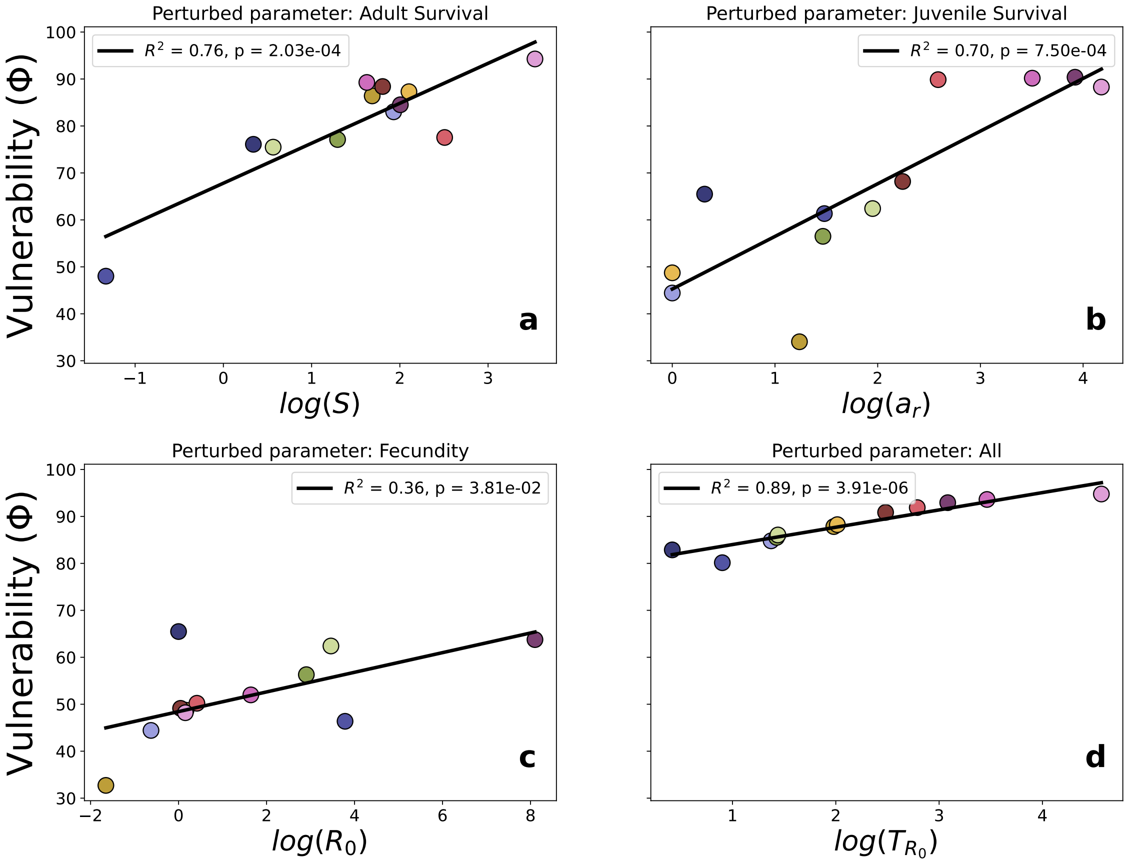


**Figure 3: Best single life-history predictors of vulnerability differ among perturbed vital rates.** Scatter plots show the strongest univariate relationship between the vulnerability metric ($\Phi$) and life-history descriptors for perturbations affecting (a) adult survival, (b) juvenile survival, (c) fecundity, and (d) all perturbed parameters combined. The best predictors were $\log(S)$, $\log(a_r)$, $\log(R_0)$, and $\log(T_{R_0})$, respectively. Black lines show linear regressions, with $R^2$ and $p$-values reported in each panel. The figure highlights that the life-history dimension most strongly associated with vulnerability depends on which vital rate is perturbed, whereas aggregate vulnerability is best captured by generation time.

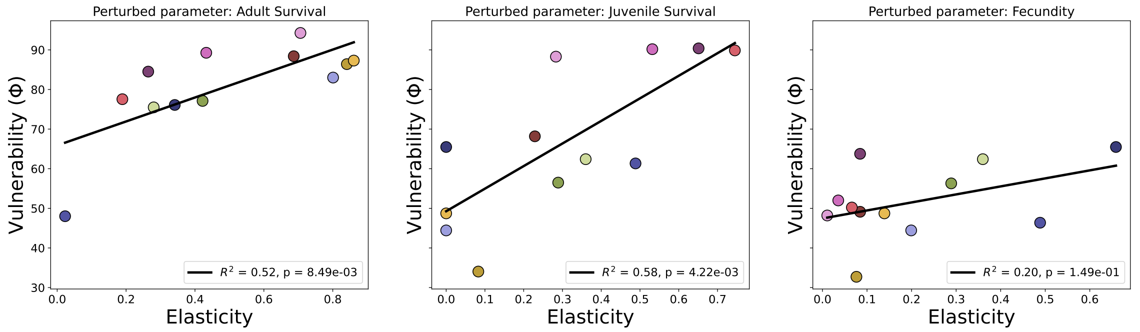


**Figure 4: Relationship between vulnerability and elasticity for each perturbed demographic parameter.** Scatter plots show the relationship between the vulnerability metric ($\Phi$) and the summed elasticity of the corresponding perturbed parameter across species for perturbations affecting adult survival, juvenile survival, and fecundity. Lines indicate linear regressions, with $R^2$ and $p$-values shown in each panel. Elasticity was positively associated with vulnerability for perturbations affecting survival, but explained only part of its variation, whereas the relationship was weaker for fecundity. This indicates that, although vulnerability is related to classical asymptotic perturbation metrics, it also captures additional information associated with the magnitude and temporal structure of perturbation regimes.
